# Species interactions are internally constrained despite large climatic variability

**DOI:** 10.1101/2025.10.28.684754

**Authors:** Sergio Picó-Jordá, Chuliang Song, Oscar Godoy

## Abstract

Understanding how vital rates and species interactions vary over time is crucial for predicting community responses to environmental change. Considerable progress has been made in understanding the drivers of variation in vital rates. However, the question of whether interactions are highly plastic and context-dependent, or strongly constrained by internal (e.g., species traits and composition) and/or external factors (e.g., environmental conditions) remains unclear. We applied a theoretical approach based on the feasibility domain —the range of conditions allowing coexistence— to a nine-year dataset of time-changing interactions between annual plants under large variability in annual precipitation. Using subcommunities of three species, we found that species interactions are strongly constrained, forming a “core-periphery” structure of consistently feasible combinations across years. This main finding means that species sample repeatedly a restricted range of opportunities for coexistence. Similar findings were obtained for subcommunities of four species. Crucially, the constraints to variation in biotic interactions are determined by species identity (internal constraints) rather than precipitation or temporal autocorrelation (external environmental factors). Furthermore, we found a contrasting effect of precipitation on the feasibility of subcommunities. While wetter years increase similarity between subcommunities and reduce the overall feasible range, drier years increase dissimilarity between subcommunities and increase the probability of coexistence when the conditions seem harsher. These findings suggest that constraints to biotic interactions tend to be alike across species in wetter years, but more context dependency occurs across species in drier years. Our findings challenge the assumption of highly plastic species interactions even in a highly dynamic system of annual plants. Our results also highlight the critical importance of internal constraints generated by species identity in mediating community persistence and predicting community responses to environmental change.

## Introduction

To predict how ecological communities will respond to a changing world, we need to understand two intertwined processes: the vital rates of individual species [Bernhardt et al., 2020, Boyce et al., 2006], and the complex web of interactions between them [Brooker, 2006, Tylianakis et al., 2008]. Organisms of the same species share characteristic traits like body size, feeding strategy, or metabolic rates that produce vital rates (survival, growth, and reproduction) that differ between species [Adler et al., 2014]. Analogously, the characteristic traits of a species determine how often it interacts with other species, the kind of interaction, how strong these interactions are, and how symmetric [Pérez-Ramos et al., 2019, Strydom et al., 2021]. Together, these two components determine population dynamics while also potentially providing species with flexibility to cope with novel fast-changing environments. While considerable progress has been made in understanding the responses of individual species vital rates [Compagnoni et al., 2021, Slein et al., 2023], a critical piece of the puzzle remains missing. Whether and how these interactions themselves —the architects of ecosystem stability— shift and reorganize in response to environmental variability remains largely unknown. Obtaining this information is critical to understand how much environmental variation a community can withstand before losing species [Grilli et al., 2017], since species interactions determine what combinations of vital rates allow for the persistence of all species in the community [Saavedra et al., 2017].

Species interactions, as a measure of the effect of one species on the growth rate of another, are not fixed. Their strength (weak vs. strong) and sign (positive vs. negative) vary markedly across space and time [CaraDonna et al., 2017, Ushio et al., 2018, Zvereva and Kozlov, 2021]. Although some species traits or abiotic conditions seem to generate some patterns [Daniel et al., 2024, Maestre and Cortina, 2004], the prevailing view holds that most changes in species interactions are highly context-dependent and hence difficult to predict [Catford et al., 2022, Chamberlain et al., 2014, Song et al., 2020]. This inherent variability and unpredictability raise a fundamental question: Are changes in species interactions essentially random, or are they constrained by underlying factors? These constraints could arise from either internal community characteristics such as species identity with their particular trait profile and evolutionary origin, or from external environmental drivers such as rainfall variability. Understanding the nature and strength of these constraints is essential for predicting how ecological communities will respond to ongoing environmental change and to develop effective conservation strategies [Tylianakis et al., 2008].

To address the dichotomy of the internal versus external forces shaping species interactions, we adopt a structuralist approach [Saavedra et al., 2017, Svirezhev and Logofet, 1978] that allows scaling up from prior pairwise approaches [Hallett et al., 2019, Kraft et al., 2015a] to consider a community-level perspective. Within this approach, a key concept is the feasibility domain, which allows quantifying the various constraints on species interactions. As an analogy, think of the feasibility domain as a “safe operating space” for the community: it represents the range of conditions —such as combinations of growth rates— where all species can coexist [Godoy et al., 2018, Saavedra et al., 2017]. A larger and more symmetric feasibility domain implies a higher probability of long-term community persistence [Allen-Perkins et al., 2023, Song et al., 2018]. While this framework has typically been used to analyze communities with fixed interactions [Saavedra et al., 2017], we extend it to a dynamic system with changing interactions: how the feasibility domain —a direct reflection of the current interaction network— itself moves and morphs in response to environmental change [Song et al., 2018, 2020].

The dynamic feasibility domain approach allows us to describe how the feasibility domains of different years explore the range of possible parameters — growth rates in this case — and if their movements between consecutive years are gradual or abrupt. Using this technique, we can test three distinct hypotheses about the nature of constraints on species interactions (Figure 1). First, strong internal constraints may dictate interactions, driven primarily by fixed species traits (e.g., phenology, functional traits) [Daniel et al., 2024, Olesen et al., 2011] or taxonomic identity [Godoy et al., 2014]. If this is the case, feasibility domains from different years would consistently overlap, forming a *core-periphery structure*, where a central region of the interaction space is consistently feasible (Figure 1 a). Second, multiple external constraints could drive shifts between distinct states of species interactions. For example, alternating periods of drought and flooding might lead to different community configurations or changes in relative abundances [Fujita et al., 2023] and, consequently, feasibility domains using distinct areas of the parameter space (Figure 1 b). Third, a lack of strong constraints might result in feasibility domains that vary randomly across the parameter space, reflecting a high degree of environmental forcing or stochasticity (e.g., strong effect of precipitation or dispersion) in community assembly (Figure 1 c). As each scenario provides different distributions of overlaps between feasibility domains of consecutive years (Figure 1 d-f), our theoretical framework provides a powerful method to distinguish between internal and external constraints on species interactions.

**Figure 1.**
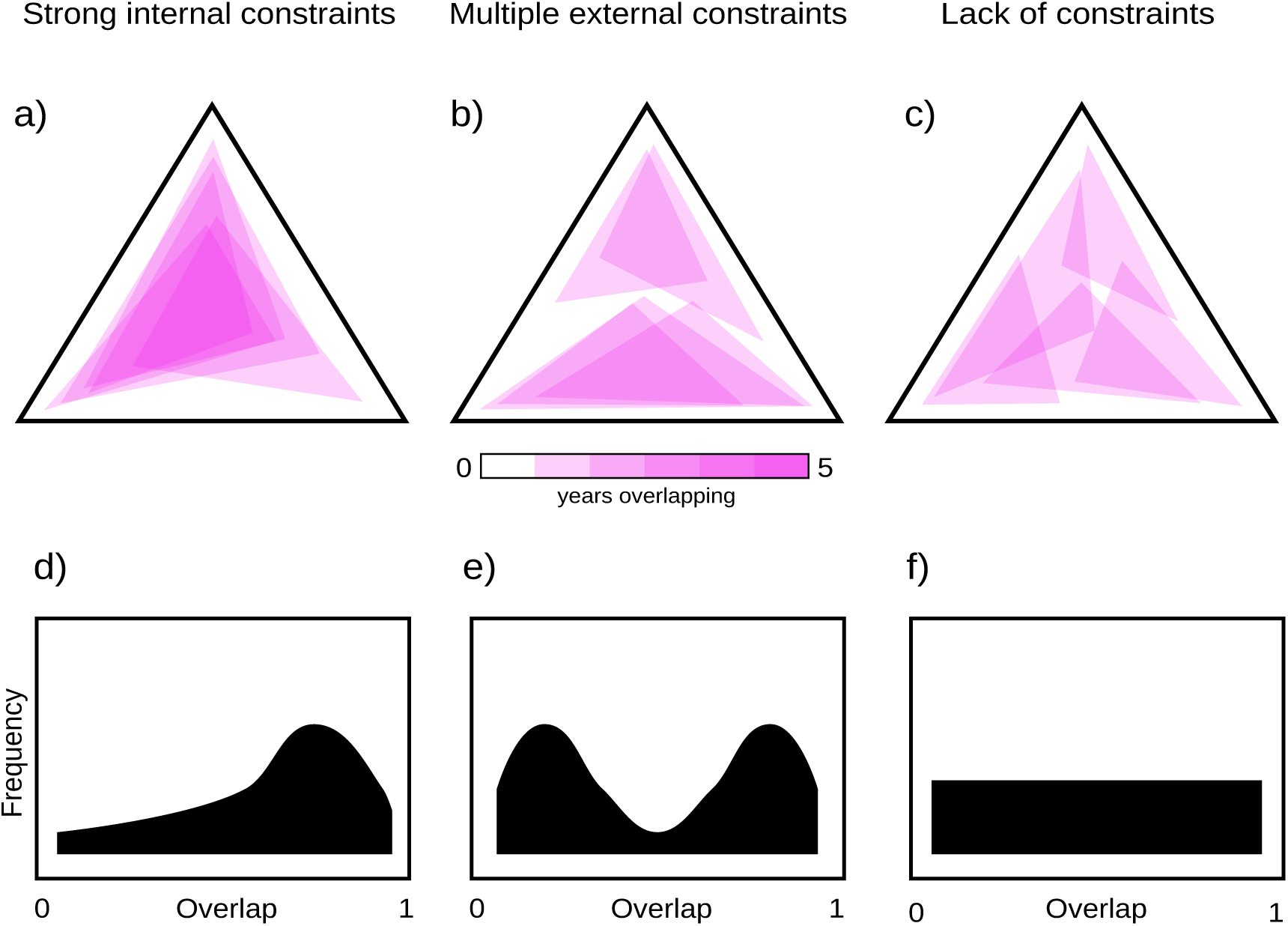
Theoretical expectations for how the feasibility domains (FDs) of a community explore the parameter space over time. (a) Strong internal constraints scenario: FDs remain concentrated in a limited region, indicating strong internal constraints. (b) Multiple external constraints scenario: FDs shift between distinct regions, suggesting multiple stable states. (c) Lack of constraints scenario: FDs spread across the entire parameter space, implying minimal constraints. (d-f) Expected distributions of overlap values between pairs of consecutive FDs for each scenario: (d) High internal constraints lead to consistently high overlap, (e) alternative stable regions result in a bimodal distribution of overlap values, and (f) no constraints produce a uniform overlap distribution.

With our approach, we hypothesize that if internal constraints are dominant, randomizing species identities within the interaction matrix should disrupt the core-periphery pattern and the observed temporal conservation of the feasibility domain. In contrast, external constraints creating a similar *core-periphery structure* act through autocorrelation in environmental conditions or by species-mediated modifications of the microenvironment. For instance, species can gradually build up thick litter layers (e.g., grasses with litter hard to decompose) or increase the abundance of natural enemies that persist in the soil (e.g., forbs with large and soft leaves that are attacked by fungi) [Bever, 2003, Bever et al., 2015]. If these externally driven, time-dependent processes are the main constraint, then the order in which species interactions occur over time becomes crucial. Specifically, interaction matrices from consecutive years should be more similar than those from more distant years. Therefore, randomizing the temporal order of these matrices should eliminate this time-dependent similarity (temporal autocorrelation) and allow an assessment of origin of the constraints.

We test these hypotheses using a unique, nine-year dataset of temporal variation in species interactions in a Mediterranean grassland community (Doñana National Park, Spain). This system is characterized by annual plants from diverse taxonomic groups, experiencing highly variable annual precipitation (Appendix S1: Figure S4), making it an ideal system for investigating to what extent internal constraints versus environmental variability shape species interactions. By integrating detailed field observations with our structuralist approach, we provide the first empirical assessment of the constraints governing species interaction variation in a multi-species natural community.

## Methods

### Study system

We collected data from our field site in Caracoles Ranch, a natural grassland with no cattle present, located within Doñana National Park (SW Spain 37.07*^◦^* N, 6.31*^◦^* W). The area has a Mediterranean climate with a mean temperature of 17.5*^◦^* C and a mean precipitation of 460 mm for the period 2000-2023. A small slope generates a gradient of soil salinity and humidity. Vegetation on this site is dominated by annual plant species, with perennial species barely present.

In September 2014, we established nine plots of 8.5 m x 8.5 m along this environmental gradient, divided into three blocks of three. Plots were separated by an average distance of 30 m (minimum 20 m) and blocks by an average distance of 300 m. Each plot was divided into 36 subplots of 1 x 1 m with corridors of 0.5 m in between to allow access for measurements. For nine growing seasons (2015-2023), we measured abundances of every species in every subplot. We used nine interaction matrices (one per year), previously estimated from these field observations following a methodology already used and tested [Garćıa-Callejas et al., 2021], for the seven most common species that were present every year in every plot (Appendix S1: Table S1). These matrices characterize the interaction structure of this community every year by including the per capita effects of each species on itself and on all other species.

### Feasibility domains and overlap

Our approach was to use mainly subcommunities of triplets of species as our unit of analysis, giving a total of 35 combinations of species, although we also extended our evaluation’s main findings to subcommunities of four species (Appendix S1: Figure S3). We chose these levels of richness because, while offering some of the complexity of multispecies systems beyond simple pairs, they produce feasibility domains big enough to create patterns of overlap and, in addition, the three species case can be graphically represented in two dimensions for an easier interpretation. Note that this approach works for combinations of three and four species from the total of seven species with available interactions data. But, as the size of the feasibility domain depends on the number of species [Dougoud et al., 2018], we cannot approach each subcommunity to the total number of species because feasibility domains tend to become smaller and the overlap patterns eventually disappear. For those particular cases, the alternative would be to measure the distance between centroids or the nearest sides of consecutive feasibility domains. We calculated the feasibility domain for each combination of subcommunity and year from the corresponding matrix of interactions. Assuming that the population dynamics in a community can be approximated by a Lotka-Volterra system, we define as feasible equilibria of that system those where all species have positive abundances. The region of the parameter space of intrinsic growth rates that leads to feasible equilibria given an interaction matrix is known as the feasibility domain [Saavedra et al., 2017, Song et al., 2018]. We calculated the size of the feasibility domain as the normalized solid angle Ω(*A*) that is equal to the probability of sampling uniformly a vector of intrinsic growth rates on the unit sphere inside the feasibility domain [Song et al., 2018]. The normalized solid angle Ω(*A*) can be defined as

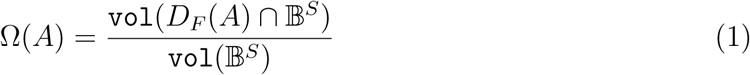

where B*^S^* is the closed unit ball in dimension S. We calculated the overlap between pairs of feasibility domains as the shared normalized solid angle Ω(*A* ∩ *B*) [Song et al., 2018], that is the range of conditions under which the community is feasible under both matrices. We calculated the overlap between pairs in three different scenarios:

1. Overlaps between feasibility domains of observed matrices in the observed order.
2. Overlaps between feasibility domains of matrices with their elements randomized in the observed order.
3. Overlaps between feasibility domains of observed matrices in a randomized order.

In our analytical design, to numerically estimate the differences in mean overlap between different scenarios with triplets, we applied a multi-membership mixed model for repeated measures to the overlap data. In that model, we incorporated the differences in mean overlap created by species identity nested in our subcommunities, since different subcommunities are not fully independent from each other, but rather they share species. We did it by implementing a presence/absence matrix of species in each observation (mean accumulated overlap for a triplet and 2-9 years) as a random factor.

### Environmental effects

To examine the effect of environmental variability, we also compiled precipitation data from the nearest meteorological station for the years 2015-2023 in the form of mm of precipitation per hydrological year (September - August) (Appendix S1: Figure S4, Estación Meteorológica de Aznalcázar, Junta de Andalućıa). For this analysis, we calculated similarity between matrices of all subcommunities in every year as the inverse of the Euclidean distance calculated as

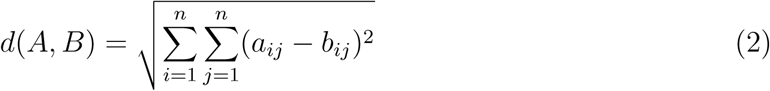

where the distance between two matrices (*A, B*) is estimated as the square root of the sum of the squares of the differences between elements *a_ij_* and *b_ij_* for every position *ij*. We also calculated mean size (Ω) of the feasibility domains in every year, and then we compared these variables against precipitation of the corresponding hydrological year (September - August). For each variable, we calculated the slope of the linear regression against precipitation with a 95% confidence interval and Spearman’s correlation.

All analyses and figures were implemented using R v4.5.0 [R Core Team, 2024] and packages *tidyverse* [Wickham et al., 2019], *feasoverlap* [Song et al., 2018], *ggtern* [Hamilton and Ferry, 2018], *lme4* [Bates et al., 2015], and *proxy* [Meyer and Buchta, 2022].

## Results

We found that species interactions are highly structured across the nine years. Rather than exhibiting random variation, we found strong evidence for internal constraints shaping the dynamics of these interactions. The feasibility domains, representing all feasible combinations of intrinsic growth rates for the case of three-species subcommunities, were not randomly distributed across the parameter space. Instead, we observed a distinct core-periphery structure (Figure 2, Appendix S1: Figure S1): certain regions of the parameter space are consistently feasible across years, forming a “core”, while other regions are only transiently feasible, constituting the “periphery”. This finding is *a priori* surprising for a strong dynamical system such as ours, where annual plant species complete their entire life cycle within a single year, and the community studied experiences strong interannual environmental variation. Thus, this persistence of the core-periphery structure strongly suggests underlying constraints that operate despite these dynamic forces.

**Figure 2.**
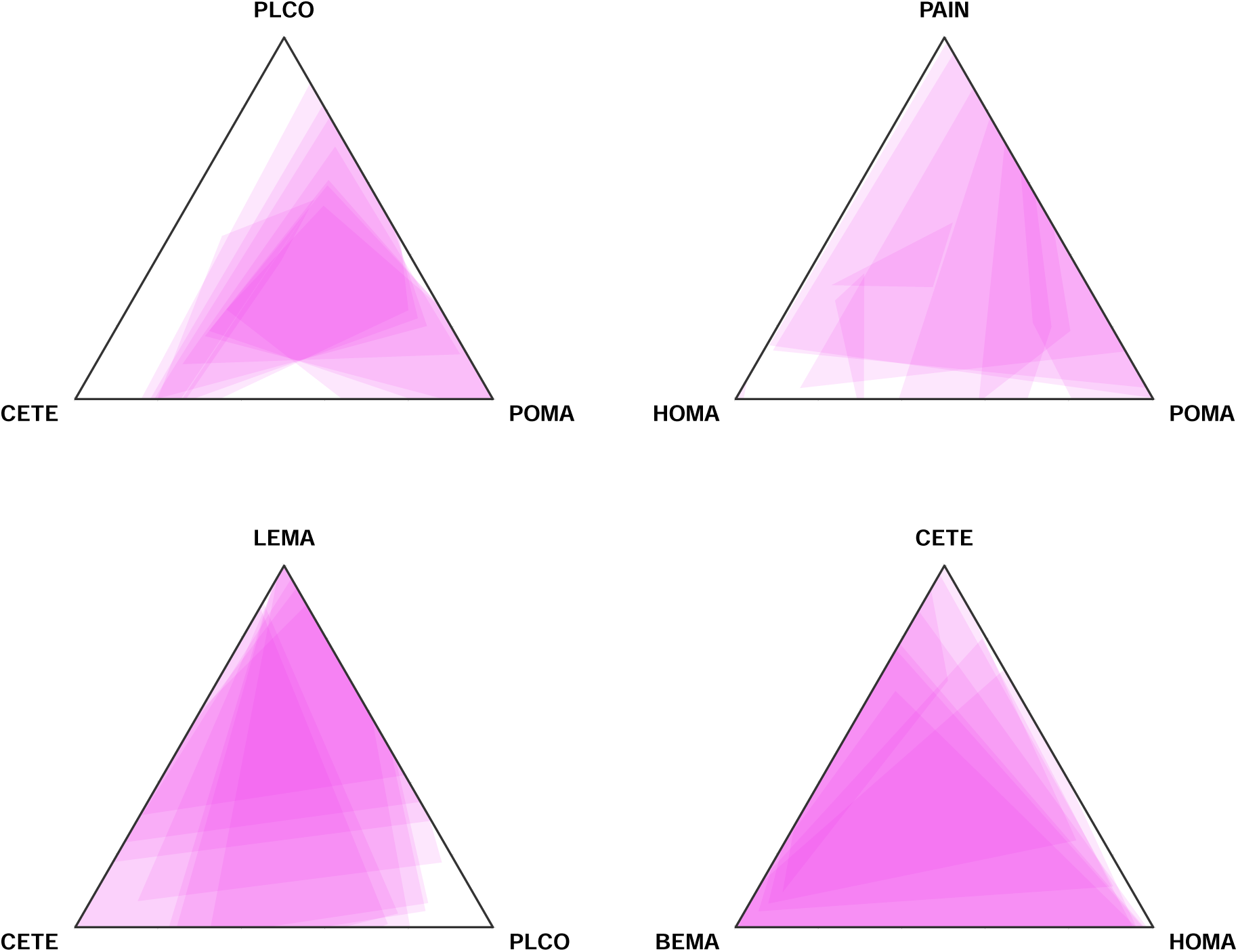
Examples of observed feasibility domains (FDs) for selected subcommunities, each composed of three species. Each pink polygon represents the FD for a given year, illustrating the range of feasible conditions for that subcommunity. Some FDs extend beyond the triangular space, indicating scenarios where at least one species has a negative intrinsic growth rate due to facilitative effects. These regions have been cropped to display only positive intrinsic growth rates. Correspondence between species codes and scientific names can be found in Appendix S1: Table S1.

To differentiate between internal and external constraints, we compared the overlap of feasibility domains between consecutive years in three scenarios: the *observed* time series, *randomized* time series (with interaction coefficients shuffled within matrices), and *disordered* time series (with the temporal order of matrices randomized) (Figure 3). This comparison rigorously accounted for repeated measurements through time and the non-independence of subcommunities (see Methods). Our results demonstrate that the degree of overlap in the observed time series significantly exceeded that of both the randomized and disordered scenarios (Figure 3 a). Notably, the largest difference was observed between the observed and randomized-interactions scenarios, with only a slight difference between the observed and disordered-time-series scenarios. These results strongly suggest that, in this system, internal constraints are the primary drivers of the core-periphery structure, with a smaller contribution from temporal autocorrelation in the structure of species interactions.

**Figure 3.**
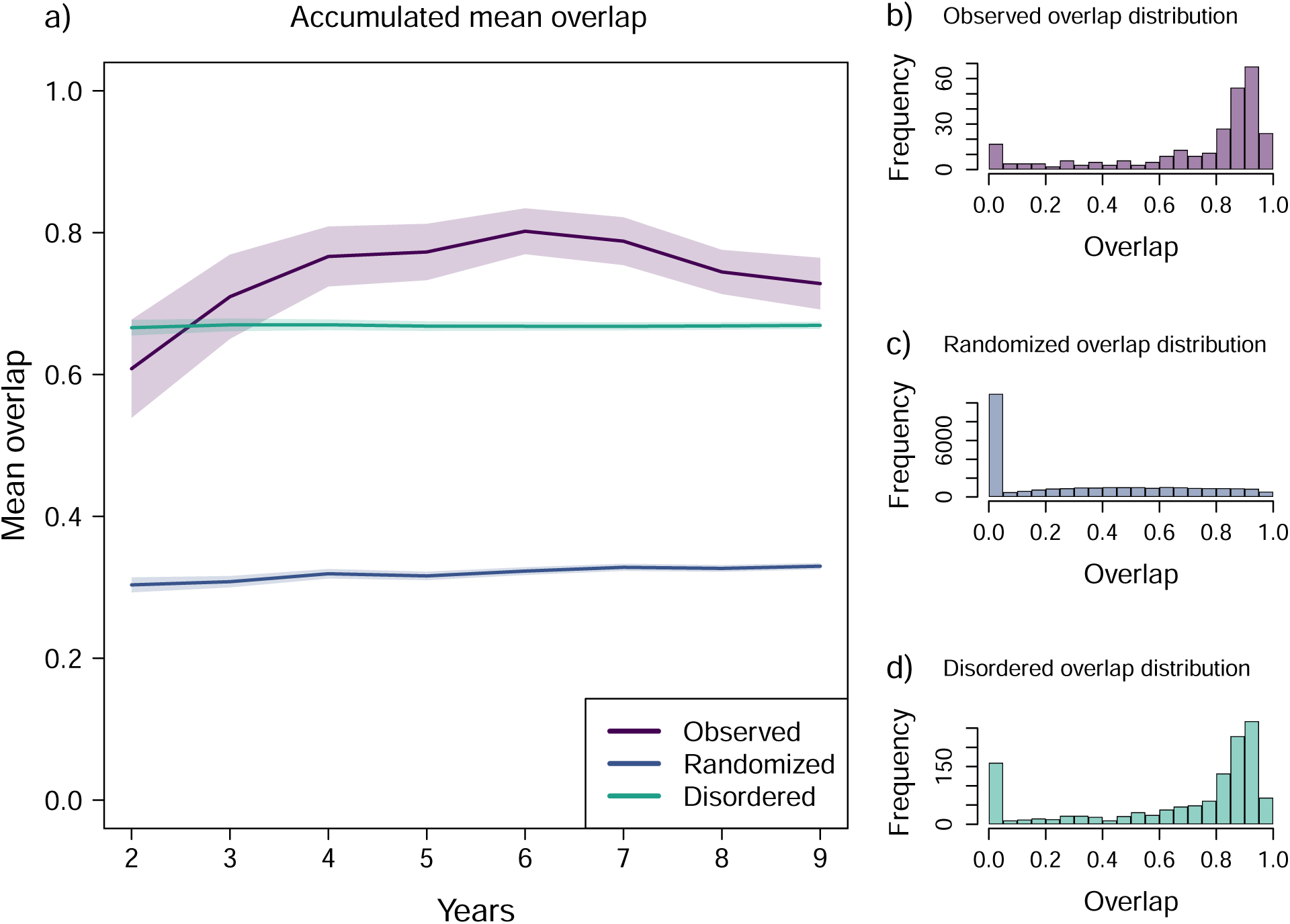
Constraints in Feasibility domains (FDs). This figure explores mean overlap between pairs of FD, calculated for subcommunities (triplets of species). Panel (a): The line plots show the accumulated mean overlap for three scenarios: (1) Observed Interactions, Observed Order (purple): Using the empirically observed matrices in their original temporal order. (2) Randomized Interactions, Observed Order (blue): Using matrices with randomized entries but maintaining the original temporal order. (3) Observed Interactions, Randomized Order (green): Using the empirically observed matrices but with their temporal order randomized. Lines represent the mean calculated across all subcommunities (triplets of species), shaded areas 95% confidence intervals for each mean. Panels (b)-(d): Distribution of Pairwise Overlap. Histograms showing the distribution of the degree of overlap between all possible pairs of matrices within each scenario. Panels display the distribution of pairwise FD overlap values for each scenario: (b) Observed, (c) Randomized Matrices, (d) Disordered.

Given the dominance of internal constraints, a remaining question is what role the environmental variability plays in our study system. We found that annual precipitation, a known driver of community dynamics in this system [Godoy et al., 2024], exerts a complex influence on the feasibility domains. Years with higher precipitation are associated with a significant reduction in the average size (Ω) of the feasibility domains across subcommunities (Figure 4 a). Simultaneously, however, the feasibility domains themselves become more similar between them in wetter years (Figure 4 b). The combination of these last two findings suggests that the wetter the conditions, the more constrained the interactions across all triplets of species to the same narrow part of the parameter space, so all subcommunities need similar conditions to coexist. Conversely, these results also suggest that two mechanisms increase the opportunities for species to coexist when the conditions are drier. First, the feasibility domain for each subcommunity tends to be bigger, and second, the feasibility domains of different subcommunities are positioned at different locations of the parameter space, which overall increases the likelihood of the system to maintain diversity by covering a larger fraction of the parameter space.

**Figure 4.**
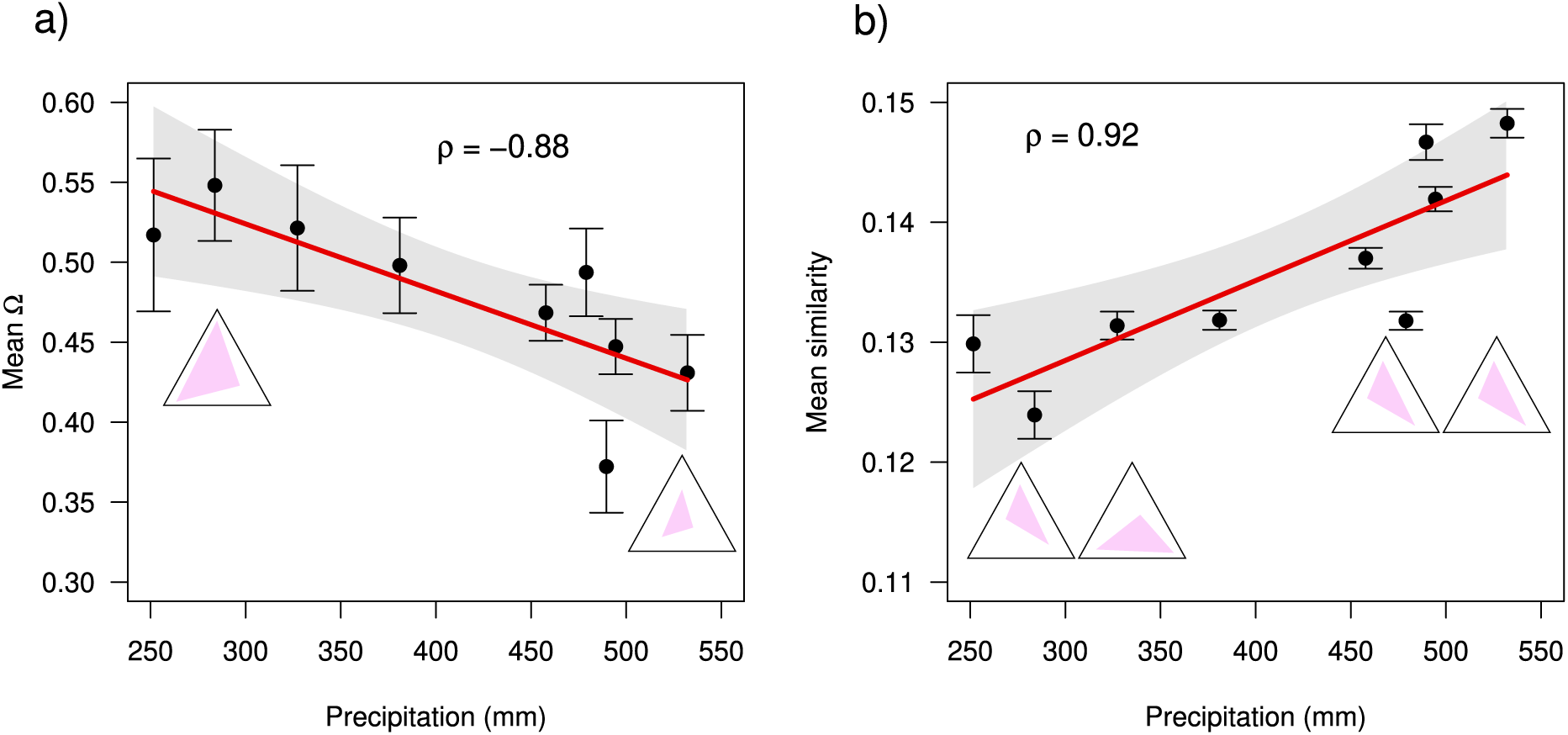
Precipitation reduces feasibility domain mean size but increases the temporal similarity of species interactions. This figure displays how annual precipitation exerts contrasting influences on two key ecological properties of subcommunities (species triplets). Panel (a) Feasibility Domain Size (Ω). The points denote the mean size (Ω) of feasibility domains, averaged across all subcommunities within a given year. Error bars represent 95% confidence intervals for these annual means. Panel (b) Interaction Matrix Similarity. The points denote the similarity between interaction matrices, quantified using the inverse of the euclidean distance between them, averaged across all subcommunities per year. Error bars represent 95% confidence intervals for these annual means. In both panels, red lines and shaded regions depict the predicted values and the 95% confidence intervals from the respective linear regressions, respectively. Also, Spearman’s correlation (*ρ*) is shown for each case. For illustrative purposes, we show for each panel examples of feasibility domains at the extremes of the precipitation gradient. In a), we highlight the differences in size between larger domains under drier years and smaller domains under wetter years. In b), we highlight differences in the position and shape of the feasibility domains of different subcommunities in drier years versus similar locations and shapes in wetter years.

## Discussion

It is widely acknowledged that species interactions are fundamental to maintaining biodiversity, yet our understanding of the temporal dynamics of these interactions remains surprisingly limited and not connected to how stable communities are against environmental variation. Part of this limitation is because much of the recent debate has centered on whether these changes are deterministic or stochastic [CaraDonna et al., 2017, Catford et al., 2022, Chamberlain et al., 2014, Daniel et al., 2024, Hallett et al., 2019, Ushio et al., 2018]. However, this focus often overlooks a more fundamental question: To what extent are species interactions constrained, as opposed to being highly plastic and adaptable? Our findings strongly support the view that interactions are constrained by the composition of species within communities, while external drivers such as precipitation —commonly believed to play a major role [Hallett et al., 2019, Matías et al., 2018, Van Dyke et al., 2022, Wainwright et al., 2019]— have a comparatively minor effect on generating variation in interactions. Additionally, we found evidence that our communities show a temporally auto-correlated structure. This finding is surprising in a community of annual plants, indicating that the system retains a “memory” of past interactions and abundances that influences short-term dynamics. While the underlying mechanisms are not yet fully understood, we hypothesize that this temporal autocorrelation could reflect gradual changes in species abundances or small-scale habitat modifications such as attracting natural enemies [Song et al., 2021] and self-limiting processes such as litter build-up [Letts et al., 2015].

The observed constraints on species interactions have significant implications for understanding the maintenance of species diversity, particularly given the increasing environmental variability driven by human activities. Our finding that these constraints are strong and internally driven suggests that communities have a limited capacity to absorb environmental change before species extinctions occur [Saavedra et al., 2017, Song et al., 2020]. Specifically, stronger constraints mean a smaller portion of the feasibility domain is accessible, increasing the likelihood that environmental fluctuations will push the intrinsic growth rates of the system outside the bounds of species positive growth rates, which can negatively impact biodiversity because species can no longer thrive in the system [Allen-Perkins et al., 2023, Grilli et al., 2017]. Consequently, even seemingly minor environmental shifts can trigger substantial changes in community composition [Van Dyke et al., 2022]. Therefore, the maintenance of biodiversity under such changing conditions relies heavily on compositional shifts: different species assemblages occupying distinct regions of the parameter space (see Appendix S1: Figure S1). This perspective offers a crucial complement to existing explanations for the widely documented pattern of species turnover across broad temporal and spatial climatic gradients [Buckley and Jetz, 2008, Korhonen et al., 2010]. While it is well-established that species adapt to local conditions —a process where the environment effectively filters the community members [HilleRisLambers et al., 2012, Kraft et al., 2015b]— our findings highlight that the inherent constraints on how species can interact within the available species pool are equally important in determining which species combinations are viable.

A key conclusion from our study is that the strength and sign of species interactions within ecological communities does not follow a completely random structure. In other words, species interactions are demonstrably constrained by biological factors, meaning that communities explore only a limited portion of the theoretically available feasibility domain. This finding has important implications for theoretical ecology, particularly for a large and influential body of work that uses network approaches based on random matrices to study species coexistence in species-rich communities [Akjouj et al., 2024, Allesina and Tang, 2012, Gibbs et al., 2022, May, 1972]. These models assume that species interactions are assigned randomly —that is, species interact with a given probability and with a strength drawn from a statistical distribution—. While we fully acknowledge the significant contributions of this random-matrix approach, our results suggest that its direct applicability to real-world communities may be limited by its inherent assumption of random interactions. A more realistic and potentially fruitful direction for future theoretical work could involve incorporating internal constraints on interactions in these network models. That is, combining a probabilistic approach [Strydom et al., 2021] with the creation of species “identities”, a set of constraints that define how often a species interacts with other species, the kind of interactions, how strong these interactions are, and how symmetric.

An intriguing finding of our study is that environmental variation in the form of precipitation variability does not have uniform effects on different facets of species interactions and feasibility. Specifically, the drier and wetter ends of the annual precipitation gradient have contrasting impacts. While drier years allow subcommunities to explore different subsections of the parameter space that are bigger in average, wetter years reduce this variation, making all subcommunities behave more alike and making their feasibility domains shrink in average. From our experience observing the study system during a decade, we deduce that these patterns emerge because in drier years competition is relaxed and may even shift to facilitation as total biomass remains low, while in wetter years, total biomass is way higher and there are enough plants and of a big enough size as to compete for resources like nutrients and light. The ecological meaning is that drier years increase the opportunities for coexistence, while the opposite is true in wetter years. These temporal processes affecting the feasibility of natural communities have never been reported before beyond our study system, and they imply that the directionality of the environmental change matters for constraining interactions and maintaining diversity.

In conclusion, our study demonstrates that even in a highly dynamic system of short-lived annual plants, species interactions are strongly constrained, primarily by the identity of the interacting species. It is reasonable to hypothesize that longer-lived organisms, such as perennial plants or trees, might exhibit even stronger constraints due to their slower growing strategies, though this requires further validation. Our study has therefore implications for how the local species pool can cope with changing environmental conditions, and makes complementary explanations for changes in species composition across broad environmental gradients. Taken together, these results underscore the critical need to incorporate the constrained nature of species interaction variability into both our understanding and our predictions of species coexistence and biodiversity maintenance.

## Acknowledgements

SP acknowledges financial support provided by the University of Cádiz (UCA/R93REC/2019). OG acknowledges financial support provided by the Spanish Ministry of Economy and Competitiveness (MINECO) and by the European Social Fund through TASTE (PID2021-127607OB-I00) project.

## Author Contributions

SP, CS, and OG conceptualized the idea, SP and CS conducted main statistical analyses, SP wrote the first version of the manuscript, CS and OG contributed significantly to editing and revision.

## Appendix S1 for

### A List of species and codes

**Table S1:**
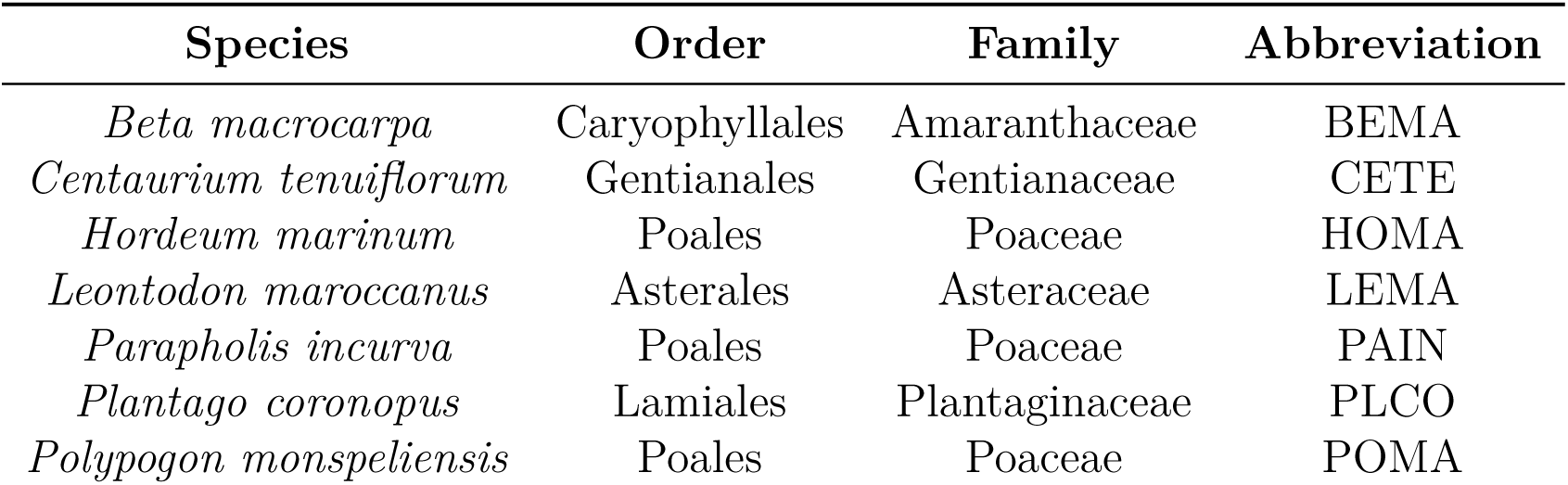
List of species used to estimate interactions and abbreviations used to identify them in code and plots.

| Species | Order | Family | Abbreviation |
| --- | --- | --- | --- |
| *Beta macrocarpa* | Caryophyllales | Amaranthaceae | BEMA |
| *Centaureum tenuiflorum* | Gentianales | Gentianaceae | CETE |
| *Hordeum marinum* | Poales | Poaceae | HOMA |
| *Leontodon maroccanus* | Asterales | Asteraceae | LEMA |
| *Parapholis incurva* | Poales | Poaceae | PAIN |
| *Plantago coronopus* | Lamiales | Plantaginaceae | PLCO |
| *Polypogon monspeliensis* | Poales | Poaceae | POMA |

### B Feasibility domains and overlap

#### B.1 Three species

**Figure S1:**
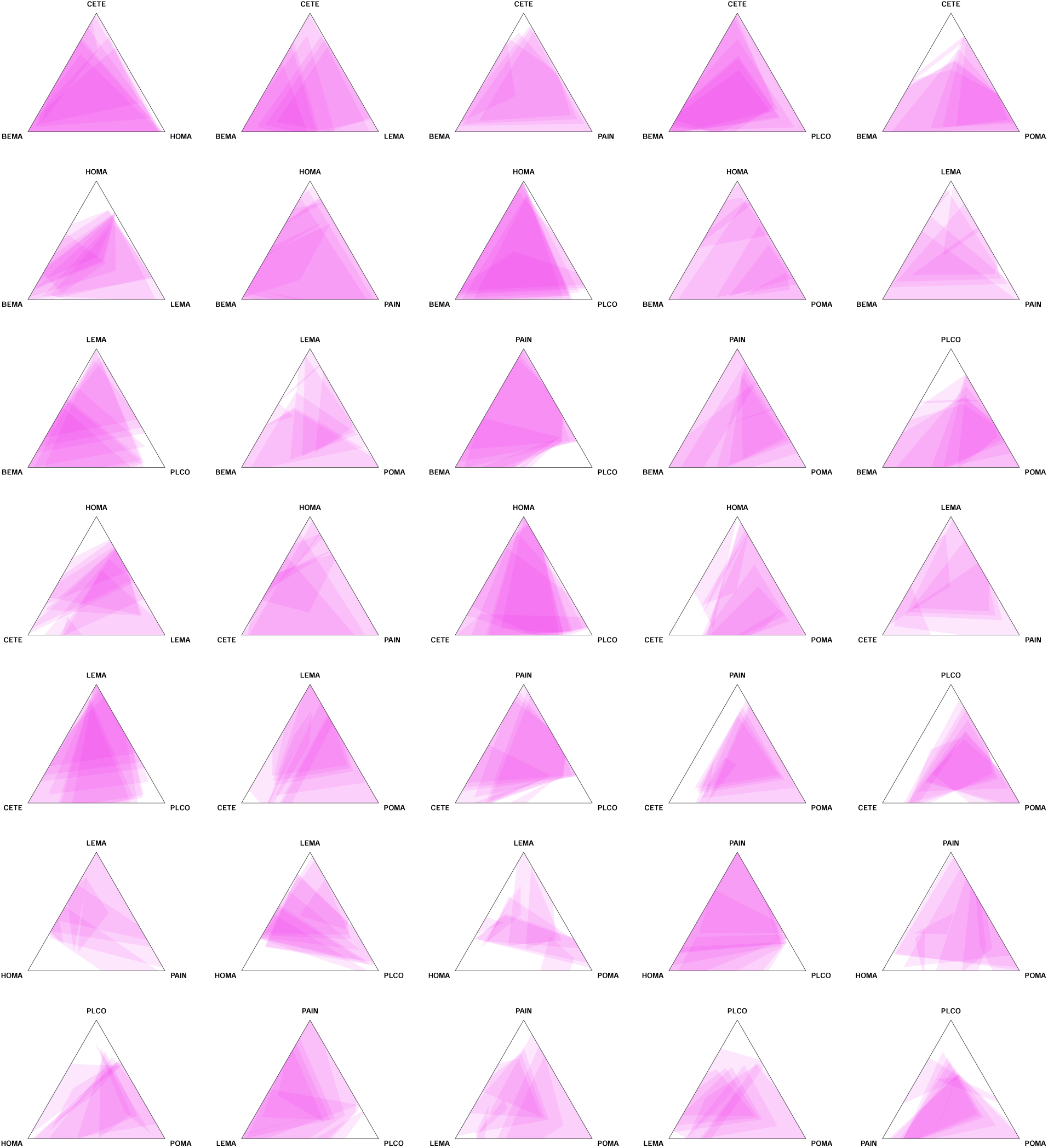
Feasibility domains (FDs) observed for all subcommunities (triplets of species). Each black line triangle contains the FDs observed during 9 years for a subcommunity. Each pink triangle represents the FD of one year for that subcommunity. Some FDs extend outside the triangle space (where the intrinsic growth rate for at least one species is negative) because of facilitative effects, but these regions have been cut to only show positive intrinsic growth rates. Correspondence between species codes and scientific names can be found in Appendix S1: Table S1.

**Figure S2:**
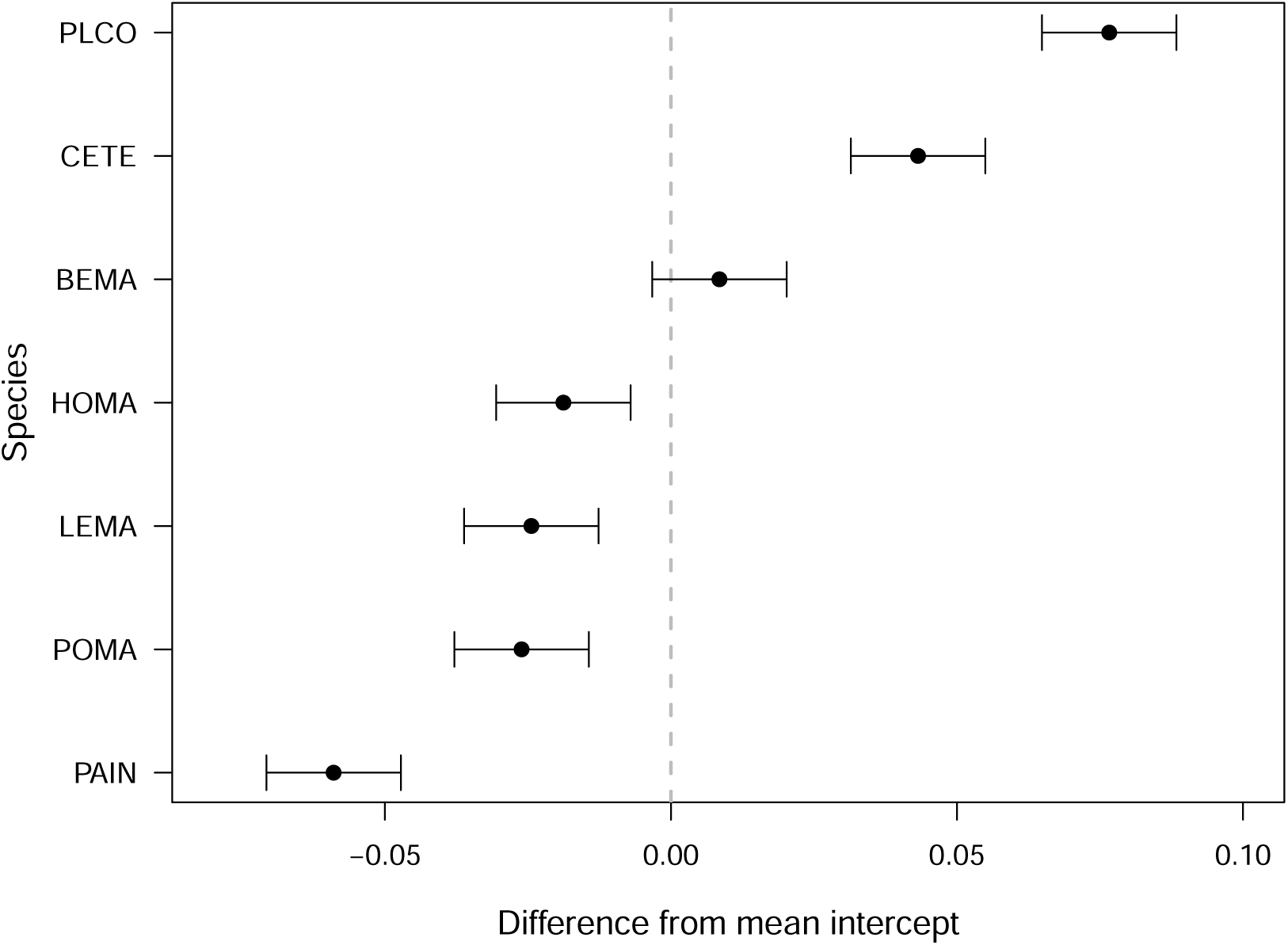
Species effects. Differences from the mean intercept for accumulated mean overlap caused by the presence of a species in a subcommunity (triplet of species). Estimated using a multimembership mixed model where treatment (Observed, Randomized, Disordered) and time (years) explain accumulated mean overlap, using a species-presence matrix as a random effect instead. Point estimates and 95% confidence intervals are shown.

**Table S2:** Multimembership model fixed effects. Summary table of the fixed effects estimated in the multimembership mixed model where treatment (Observed, Randomized, Disordered) and time (years) explain accumulated mean overlap, and a species-presence matrix is used as a random effect instead.

| **Predictor** | **Estimate** | **Std. error** | **t value** |
| --- | --- | --- | --- |
| Disordered (Intercept) | 0.642 | 0.052 | 12.377 |
| Observed | 0.067 | 0.008 | 8.221 |
| Randomized | -0.350 | 0.008 | -42.712 |
| Time (years) | 0.006 | 0.001 | 3.781 |

#### B.2 Four species

**Figure S3:**
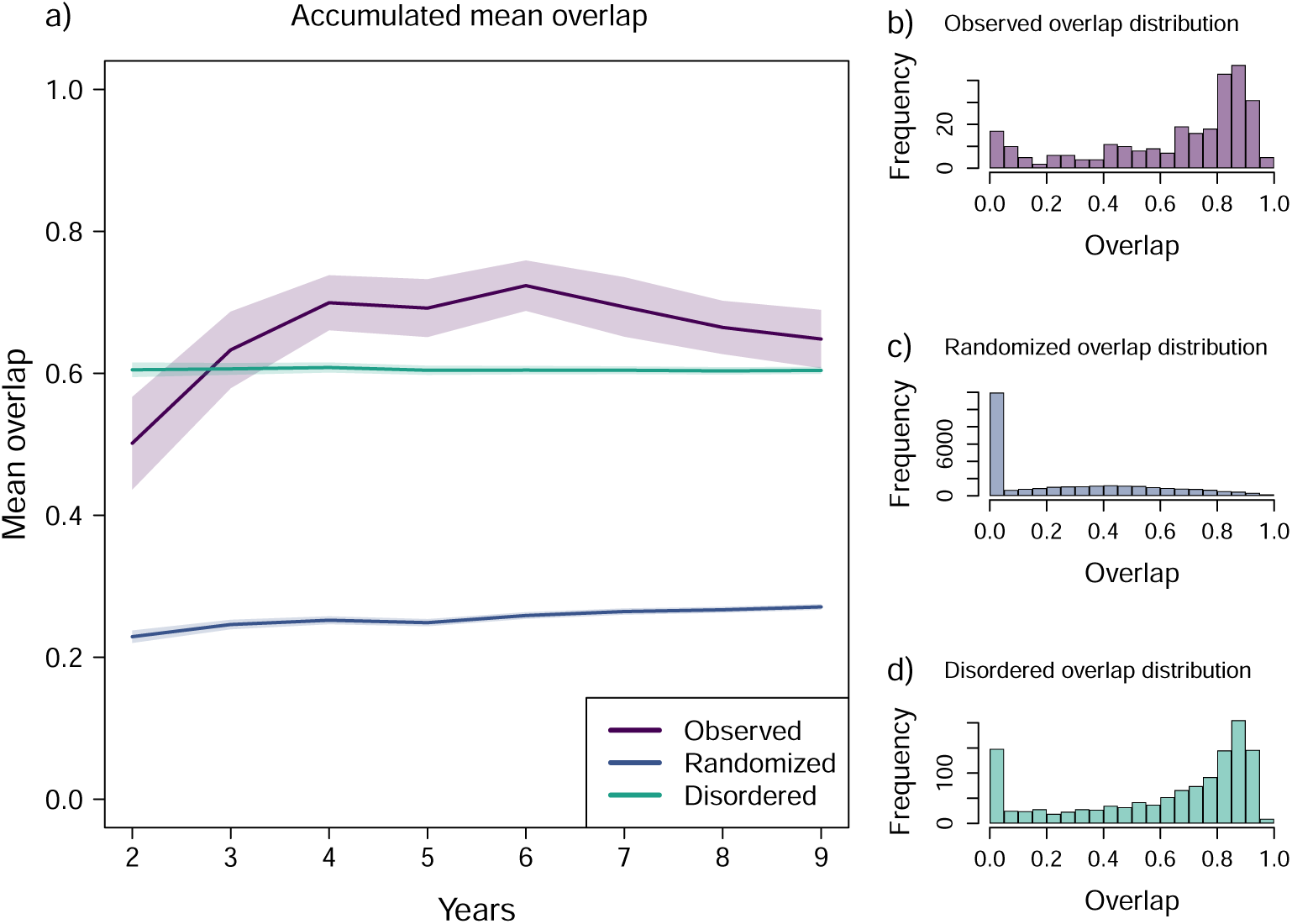
Constraints in Feasibility domains (FDs). This figure explores mean overlap between pairs of FD, calculated for subcommunities (groups of four of species). Panel (a): The line plots show the accumulated mean overlap for three scenarios: (1) Observed Interactions, Observed Order (purple): Using the empirically observed matrices in their original temporal order. (2) Randomized Interactions, Observed Order (blue): Using matrices with randomized entries but maintaining the original temporal order. (3) Observed Interactions, Randomized Order (green): Using the empirically observed matrices but with their temporal order randomized. Lines represent the mean calculated across all subcommunities (groups of four of species), shaded areas 95% confidence intervals for each mean. Panels (b)-(d): Distribution of Pairwise Overlap. Histograms showing the distribution of the degree of overlap between all possible pairs of matrices within each scenario. Panels display the distribution of pairwise FD overlap values for each scenario: (b) Observed, (c) Randomized Matrices, (d) Disordered.

### C Precipitation and precipitation effects

**Figure S4:**
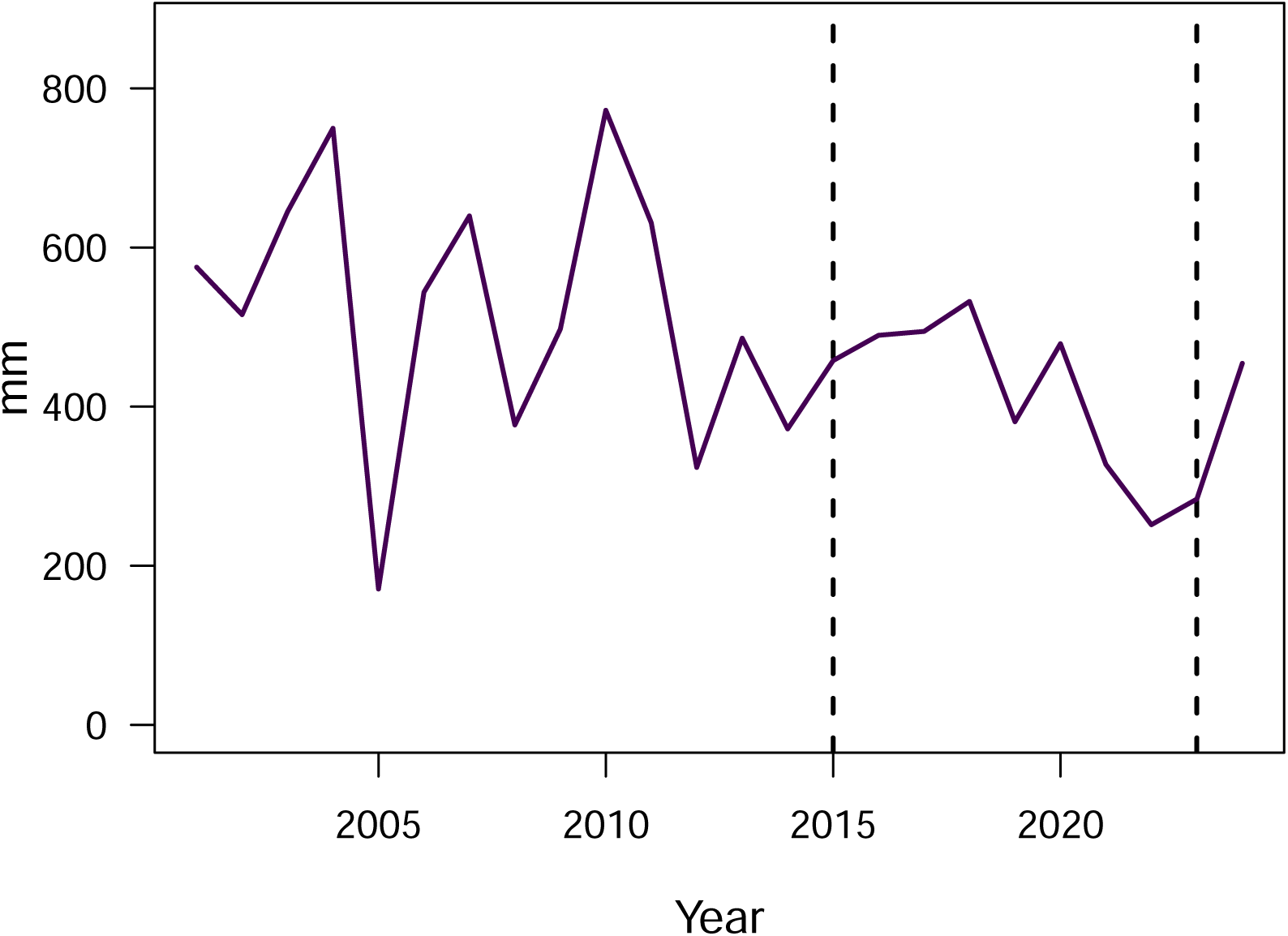
Precipitation time series from the study site. Precipitation data obtained from the nearest meteorological station to the study site (∼10 km), Estación Meteorológica de Aznalcázar, Junta de Andalucía. Data is expressed as total mm of precipitation for the hydrological year (September - August). Vertical dashed bars mark the duration of the study (2015 - 2023).

**Table S3:** Summary table of the linear regression shown in Figure 4 a).

| **Predictor** | **Estimate** | **Std. error** | **p-value** |
| --- | --- | --- | --- |
| Intercept | 0.6500 | 0.0508 | 4.1500E-06 |
| Precipitation | -0.0004 | 0.0001 | 0.0102 |

**Table S4:**
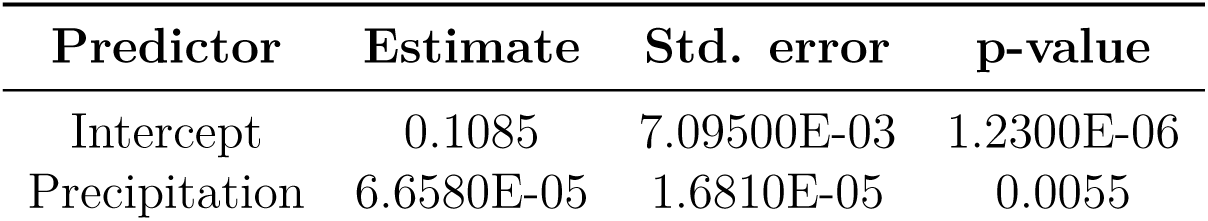
Summary table of the linear regression shown in Figure 4 b).

| **Predictor** | **Estimate** | **Std. error** | **p-value** |
| --- | --- | --- | --- |
| Intercept | 0.1085 | 7.09500E-03 | 1.2300E-06 |
| Precipitation | 6.6580E-05 | 1.6810E-05 | 0.0055 |

## References

Peter B. Adler, Roberto Salguero-Gómez, Aldo Compagnoni, Joanna S. Hsu, Jayanti Ray-Mukherjee, Cyril Mbeau-Ache, and Miguel Franco. Functional traits explain variation in plant life history strategies. Proceedings of the National Academy of Sciences, 111(2):740–745, 2014. doi: 10.1073/pnas.1315179111.

Imane Akjouj, Matthieu Barbier, Maxime Clenet, Walid Hachem, Mylène Maïda, François Massol, Jamal Najim, and Viet Chi Tran. Complex systems in ecology: a guided tour with large Lotka–Volterra models and random matrices. Proceedings of the Royal Society A: Mathematical, Physical and Engineering Sciences, 480(2285):20230284, 2024. doi: 10.1098/rspa.2023.0284.

Alfonso Allen-Perkins, David García-Callejas, Ignasi Bartomeus, and Oscar Godoy. Structural asymmetry in biotic interactions as a tool to understand and predict ecological persistence. Ecology Letters, 26(10):1647–1662, 2023. doi: 10.1111/ele.14291.

Stefano Allesina and Si Tang. Stability Criteria for Complex Ecosystems. Nature, 483 (7388):205–208, 2012. doi: 10.1038/nature10832.

Douglas Bates, Martin Mächler, Ben Bolker, and Steve Walker. Fitting linear mixed-effects models using lme4. Journal of Statistical Software, 67(1):1–48, 2015. doi: 10.18637/jss.v067.i01.

Joey R. Bernhardt, Mary I. O’Connor, Jennifer M. Sunday, and Andrew Gonzalez. Life in fluctuating environments. Philosophical Transactions of the Royal Society B: Biological Sciences, 375(1814):20190454, 2020. doi: 10.1098/rstb.2019.0454.

James D. Bever. Soil community feedback and the coexistence of competitors: conceptual frameworks and empirical tests. New Phytologist, 157(3):465–473, 2003. doi: 10.1046/j.1469-8137.2003.00714.x.

James D. Bever, Scott A. Mangan, and Helen M. Alexander. Maintenance of Plant Species Diversity by Pathogens. *Annual Review of Ecology*, Evolution, and Systematics, 46(1): 305–325, 2015. doi: 10.1146/annurev-ecolsys-112414-054306.

Mark Boyce, Chirakkal V. Haridas, Charlotte T. Lee, and NCEAS Stochastic Demosgraphy Working Group. Demography in an increasingly variable world. Trends in Ecology & Evolution, 21(3):141–148, 2006. doi: 10.1016/j.tree.2005.11.018.

Rob W. Brooker. Plant–plant interactions and environmental change. New Phytologist, 171 (2):271–284, 2006. doi: 10.1111/j.1469-8137.2006.01752.x.

Lauren B. Buckley and Walter Jetz. Linking global turnover of species and environments. Proceedings of the National Academy of Sciences, 105(46):17836–17841, 2008. doi: 10.1073/pnas.0803524105.

Paul J. CaraDonna, William K. Petry, Ross M. Brennan, James L. Cunningham, Judith L. Bronstein, Nickolas M. Waser, and Nathan J. Sanders. Interaction rewiring and the rapid turnover of plant–pollinator networks. Ecology Letters, 20(3):385–394, 2017. doi: 10.1111/ele.12740.

Jane A. Catford, John R.U. Wilson, Petr Pyšek, Philip E. Hulme, and Richard P. Duncan. Addressing context dependence in ecology. Trends in Ecology & Evolution, 37(2): 158–170, 2022. doi: 10.1016/j.tree.2021.09.007.

Scott A. Chamberlain, Judith L. Bronstein, and Jennifer A. Rudgers. How context dependent are species interactions? Ecology Letters, 17(7):881–890, 2014. doi: 10.1111/ele.12279.

Aldo Compagnoni, Sam Levin, Dylan Z. Childs, Stan Harpole, Maria Paniw, Gesa Römer, Jean H. Burns, Judy Che-Castaldo, Nadja Rüger, Georges Kunstler, Joanne M. Bennett, C. Ruth Archer, Owen R. Jones, Roberto Salguero-Gómez, and Tiffany M. Knight. Herbaceous perennial plants with short generation time have stronger responses to climate anomalies than those with longer generation time. Nature Communications, 12 (1):1824, 2021. doi: 10.1038/s41467-021-21977-9.

Caroline Daniel, Eric Allan, Hugo Saiz, and Oscar Godoy. Fast–slow traits predict competition network structure and its response to resources and enemies. Ecology Letters, 27(4):e14425, 2024. doi: 10.1111/ele.14425.

Michaël Dougoud, Laura Vinckenbosch, Rudolf P. Rohr, Louis-Félix Bersier, and Christian Mazza. The feasibility of equilibria in large ecosystems: A primary but neglected concept in the complexity-stability debate. PLOS Computational Biology, 14(2): e1005988, 2018. doi: 10.1371/journal.pcbi.1005988.

Hiroaki Fujita, Masayuki Ushio, Kenta Suzuki, Masato S. Abe, Masato Yamamichi, Koji Iwayama, Alberto Canarini, Ibuki Hayashi, Keitaro Fukushima, Shinji Fukuda, E. Toby Kiers, and Hirokazu Toju. Alternative stable states, nonlinear behavior, and predictability of microbiome dynamics. Microbiome, 11(1):63, 2023. doi: 10.1186/s40168-023-01474-5.

David García-Callejas, Ignasi Bartomeus, and Oscar Godoy. The spatial configuration of biotic interactions shapes coexistence-area relationships in an annual plant community. Nature Communications, 2021. doi: 10.1101/2021.04.02.438211.

Theo Gibbs, Simon A. Levin, and Jonathan M. Levine. Coexistence in diverse communities with higher-order interactions. Proceedings of the National Academy of Sciences, 119 (43):e2205063119, 2022.

Oscar Godoy, Nathan J.B. Kraft, and Jonathan M. Levine. Phylogenetic relatedness and the determinants of competitive outcomes. Ecology letters, 17(7):836–844, 2014.

Oscar Godoy, Ignasi Bartomeus, Rudolf P. Rohr, and Serguei Saavedra. Towards the integration of niche and network theories. Trends in Ecology & Evolution, 33(4):287–300, 2018.

Oscar Godoy, Fernando Soler-Toscano, José R. Portillo, and José A. Langa. The assembly and dynamics of ecological communities in an ever-changing world. Ecological Monographs, page e1633, 2024.

Jacopo Grilli, Matteo Adorisio, Samir Suweis, György Barabás, Jayanth R. Banavar, Stefano Allesina, and Amos Maritan. Feasibility and coexistence of large ecological communities. Nature Communications, 8(1):14389, 2017. doi: 10.1038/ncomms14389.

Lauren M. Hallett, Lauren G. Shoemaker, Caitlin T. White, and Katharine N. Suding. Rainfall variability maintains grass-forb species coexistence. Ecology Letters, 22(10): 1658–1667, 2019.

Nicholas E. Hamilton and Michael Ferry. ggtern: Ternary diagrams using ggplot2. *Journal of Statistical Software*, Code Snippets, 87(3):1–17, 2018. doi: 10.18637/jss.v087.c03.

Janneke HilleRisLambers, Peter B. Adler, W. Stanley Harpole, Jonathan M. Levine, and Margaret M. Mayfield. Rethinking community assembly through the lens of coexistence theory. Annual review of ecology, evolution, and systematics, 43(1):227–248, 2012.

Jenni J. Korhonen, Janne Soininen, and Helmut Hillebrand. A quantitative analysis of temporal turnover in aquatic species assemblages across ecosystems. Ecology, 91(2): 508–517, 2010. doi: 10.1890/09-0392.1.

Nathan J. B. Kraft, Oscar Godoy, and Jonathan M. Levine. Plant functional traits and the multidimensional nature of species coexistence. Proceedings of the National Academy of Sciences, 112(3):797–802, 2015a. doi: 10.1073/pnas.1413650112.

Nathan J.B. Kraft, Peter B. Adler, Oscar Godoy, Emily C. James, Steve Fuller, and Jonathan M. Levine. Community assembly, coexistence and the environmental filtering metaphor. Functional ecology, 29(5):592–599, 2015b.

Brittany Letts, Eric G. Lamb, Jenalee M. Mischkolz, and James T. Romo. Litter accumulation drives grassland plant community composition and functional diversity via leaf traits. Plant Ecology, 216(3):357–370, 2015. doi: 10.1007/s11258-014-0436-6.

Fernando T. Maestre and Jordi Cortina. Do positive interactions increase with abiotic stress? A test from a semi-arid steppe. Proceedings of the Royal Society of London. Series B: Biological Sciences, 271, 2004. doi: 10.1098/rsbl.2004.0181.

Luis Matías, Oscar Godoy, Lorena Gómez-Aparicio, and Ignacio M. Pérez-Ramos. An experimental extreme drought reduces the likelihood of species to coexist despite increasing intransitivity in competitive networks. Journal of Ecology, 106(3):826–837, 2018.

Robert M. May. Will a Large Complex System be Stable? Nature, 238(5368):413–414, 1972. doi: 10.1038/239137a0.

David Meyer and Christian Buchta. proxy: Distance and Similarity Measures, 2022. URL https://CRAN.R-project.org/package=proxy. R package version 0.4-27.

Jens M. Olesen, Jordi Bascompte, Yoko L. Dupont, Heidi Elberling, Claus Rasmussen, and Pedro Jordano. Missing and forbidden links in mutualistic networks. Proceedings of the Royal Society B: Biological Sciences, 278(1706):725–732, 2011. doi: 10.1098/rspb.2010.1371.

Ignacio M. Pérez-Ramos, Luis Matías, Lorena Gómez-Aparicio, and Óscar Godoy. Functional traits and phenotypic plasticity modulate species coexistence across contrasting climatic conditions. Nature communications, 10(1):2555, 2019.

R Core Team. R: A Language and Environment for Statistical Computing. R Foundation for Statistical Computing, Vienna, Austria, 2024. URL https://www.R-project.org/.

Serguei Saavedra, Rudolf P. Rohr, Jordi Bascompte, Oscar Godoy, Nathan J. B. Kraft, and Jonathan M. Levine. A structural approach for understanding multispecies coexistence. Ecological Monographs, 87(3):470–486, 2017. doi: 10.1002/ecm.1263.

Margaret A. Slein, Joey R. Bernhardt, Mary I. O’Connor, and Samuel B. Fey. Effects of thermal fluctuations on biological processes: a meta-analysis of experiments manipulating thermal variability. Proceedings of the Royal Society B: Biological Sciences, 290(1992):20222225, 2023. doi: 10.1098/rspb.2022.2225.

Chuliang Song, Rudolf P. Rohr, and Serguei Saavedra. A guideline to study the feasibility domain of multi-trophic and changing ecological communities. Journal of Theoretical Biology, 450:30–36, 2018. doi: 10.1016/j.jtbi.2018.04.030.

Chuliang Song, Sarah Von Ahn, Rudolf P. Rohr, and Serguei Saavedra. Towards a Probabilistic Understanding About the Context-Dependency of Species Interactions. Trends in Ecology & Evolution, page S0169534719303568, 2020. doi: 10.1016/j.tree.2019.12.011.

Xiaoyang Song, Jun Ying Lim, Jie Yang, and Matthew Scott Luskin. When do Janzen–Connell effects matter? A phylogenetic meta-analysis of conspecific negative distance and density dependence experiments. Ecology Letters, 24(3):608–620, 2021. doi: 10.1111/ele.13665.

Tanya Strydom, Michael D. Catchen, Francis Banville, Dominique Caron, Gabriel Dansereau, Philippe Desjardins-Proulx, Norma R. Forero-Muñoz, Gracielle Higino, Benjamin Mercier, Andrew Gonzalez, Dominique Gravel, Laura Pollock, and Timothée Poisot. A roadmap towards predicting species interaction networks (across space and time). Philosophical Transactions of the Royal Society B: Biological Sciences, 376(1837): 20210063, 2021. doi: 10.1098/rstb.2021.0063.

Yuri M. Svirezhev and Dimitrii O. Logofet. The stability of biological communities. Mir Publishers, Moscow, Russia, 1978.

Jason M. Tylianakis, Raphael K. Didham, Jordi Bascompte, and David A. Wardle. Global change and species interactions in terrestrial ecosystems. Ecology Letters, 11(12): 1351–1363, 2008. doi: 10.1111/j.1461-0248.2008.01250.x.

Masayuki Ushio, Chih-hao Hsieh, Reiji Masuda, Ethan R Deyle, Hao Ye, Chun-Wei Chang, George Sugihara, and Michio Kondoh. Fluctuating interaction network and time-varying stability of a natural fish community. Nature, 554(7692):360–363, 2018. doi: 10.1038/nature25504.

Mary N. Van Dyke, Jonathan M. Levine, and Nathan J.B. Kraft. Small rainfall changes drive substantial changes in plant coexistence. Nature, 611(7936):507–511, 2022.

Claire E. Wainwright, Janneke HilleRisLambers, Hao R. Lai, Xingwen Loy, and Margaret M. Mayfield. Distinct responses of niche and fitness differences to water availability underlie variable coexistence outcomes in semi-arid annual plant communities. Journal of Ecology, 107(1):293–306, 2019. doi: 10.1111/1365-2745.13056.

Hadley Wickham, Mara Averick, Jennifer Bryan, Winston Chang, Lucy D’Agostino McGowan, Romain François, Garrett Grolemund, Alex Hayes, Lionel Henry, Jim Hester, Max Kuhn, Thomas Lin Pedersen, Evan Miller, Stephan Milton Bache, Kirill Müller, Jeroen Ooms, David Robinson, Dana Paige Seidel, Vitalie Spinu, Kohske Takahashi, Davis Vaughan, Claus Wilke, Kara Woo, and Hiroaki Yutani. Welcome to the tidyverse. Journal of Open Source Software, 4(43):1686, 2019. doi: 10.21105/joss.01686.

Elena L. Zvereva and Mikhail V. Kozlov. Latitudinal gradient in the intensity of biotic interactions in terrestrial ecosystems: Sources of variation and differences from the diversity gradient revealed by meta-analysis. Ecology Letters, page ele.13851, 2021. doi: 10.1111/ele.13851.

